# Hippocampal-somatomotor coactivation and perceived benefit following Mindfulness-Based Cognitive Therapy for Life in healthcare students: An exploratory randomized study

**DOI:** 10.64898/2026.10.12.758559

**Authors:** Nicolas Clairis, Luigi Saccaro, Sylvain Delplanque, Claire Holman, Zeynep Celen, Claudia Cachet, Camille Serquet, Ben Meuleman, Guido Bondolfi, Françoise Jermann, Serge Rudaz, Camille Piguet

## Abstract

Mindfulness-based interventions are known to enhance emotion regulation and stress management, yet their neural mechanisms remain incompletely understood. Effective coping strategies for emotion regulation and stress management would be particularly beneficial for healthcare students, frequently exposed to highly stressful situations. Based on prior work implicating hippocampal-somatomotor dynamics in emotion regulation, we tested whether Mindfulness-Based Cognitive Therapy for Life (MBCT-L) modulates hippocampal micro-coactivation patterns (μCAPs) in healthcare students. In a randomized controlled study including 161 healthcare students, 57 participants (39 females; mean age: 23±5 years old) completed resting-state fMRI before and after the 8-weeks intervention and were allocated to MBCT-L (N = 24) or a no-intervention control (N = 33) condition. Hippocampus-seeded CAPs analysis identified six recurrent patterns, including a somatomotor network. MBCT-L improved self-reported mindfulness, but no significant Group × Time interaction for hippocampus-somatomotor coactivation or resting physiological measures was found. In the MBCT-L group, higher hippocampus-somatomotor co-activity was associated with higher perceived importance of the intervention, perceived long-lasting benefits, and post-intervention mindfulness scores. This study suggests that the capacity to up-regulate hippocampus-somatomotor coactivation may index part of the individual benefit derived from mindfulness, supporting its role as a tool for emotion regulation and stress management.

## Introduction

Healthcare students are exposed to considerable stress over the course of their studies (Walter *et al*., 2013; La Fauci *et al*., 2023). Finding ways and efficient interventions to reduce stress, anxiety, burnout and depression in this target population has been a challenge for many years (Johnson *et al*., 2020; Aloufi *et al*., 2021; Fitzgibbon and Murphy, 2023). Mindfulness is a promising approach to learn to better cope with stressful situations (Khoury *et al*., 2015; Sparacio *et al*., 2024). Its effectiveness has been demonstrated not only at the behavioral level, but also when looking at physiological markers of stress such as heart-rate or blood pressure which have been observed to decrease after a mindfulness intervention (Pascoe *et al*., 2017). Additionally, mindfulness has already been demonstrated to benefit healthcare students (Barbosa *et al*., 2013; Braun *et al*., 2020; La Torre *et al*., 2022; Harrison *et al*., 2024). However, the brain mechanisms involved in this process remain to be characterized using specific and replicable mindfulness practices.

Mindfulness-based Cognitive Therapy for Life (MBCT-L) is a mindfulness intervention which has been developed by the Oxford Mindfulness Foundation designed for the general population rather than for clinical groups. It integrates mindfulness practices with insights from cognitive behavioral therapies (CBT) (Bernard *et al*., 2017). Its aim is twofold: helping people to notice and respond differently to automatic patterns of reactivity in the face of everyday stress and to nurture well-being and flourishing. MBCT-L has already been shown to improve stress management in healthcare professionals of multiple health centers in England (Strauss *et al*., 2021) or Tunisia (Kckaou *et al*., 2023). However, it remains to be seen whether these results can be confirmed in different situations or countries, as an early intervention in healthcare students, and in association to markers of brain connectivity.

Indeed, previous research suggests that mindfulness practice (Sezer *et al*., 2022; Guidotti *et al*., 2023) as well as dispositional mindfulness (Celen *et al*., 2024) modulate large-scale brain connectivity, even at rest. More specifically, independent of the practice type, mindfulness induces a general decrease in the activation of the default mode network (DMN) during practice (Brewer *et al*., 2011; Ganesan *et al*., 2023), during task execution (Berkovich-Ohana *et al*., 2016; Hehr *et al*., 2022) and during resting-state (Berkovich-Ohana *et al*., 2016) in mindfulness practitioners. In parallel, research from our own research group highlights that mindfulness practice increases the coactivation pattern occurrence between the hippocampus and the somatomotor network (SMN) at rest in teenagers after a MBCT-L intervention (Saccaro *et al*., In Preparation), suggesting that brain networks other than the DMN may also be relevant to understand the health benefits driven by mindfulness.

Convergingly, recent evidence points to a role of the SMN in emotion regulation (Kropf *et al*., 2019; Zanella *et al*., 2022; Jiang *et al*., 2026). Previous work from our group has demonstrated that the hippocampus coactivation pattern occurrence with the SMN (referred to as sensorimotor) is transdiagnostically reduced in patients with emotion dysregulation disorders as well as in at-risk offspring and that lower hippocampus-SMN coactivation pattern occurrence is associated with higher emotion dysregulation (Saccaro *et al*., 2023; Saccaro, Delavari, Van De Ville, *et al*., 2025; Saccaro, Delavari, Meuleman, *et al*., 2025). This evidence suggests that this network plays an essential role in emotion regulation, that may be influenced by mindfulness practice.

Although healthcare students face sustained academic and clinical stress that raises their risk of burnout, anxiety, and depression (Di Mario *et al*., 2024), scalable preventive interventions for this population remain underused and their neural mechanisms poorly characterized. Mindfulness, and more specifically MBCT-L, is a promising candidate to enhance well-being in healthcare students as it may result in lowering stress and improving stress handling (Khoury *et al*., 2015; Pascoe *et al*., 2017; Strauss *et al*., 2021). However, whether its benefits are accompanied by measurable changes in intrinsic brain function and whether such changes index who benefits from it is currently unknown. In this study, 161 healthcare students were enrolled in a randomized controlled trial with two aims. First, at the behavioral level, we tested the hypothesis that MBCT-L, relative to a no-intervention control, improves self-reported mindfulness and the capacity to cope with stress in daily life. The full-trial behavioral analysis is beyond the scope of this article and will be reported separately (Jermann *et al*., In Preparation). Here, we report these outcomes only for the subset of participants who underwent resting-state fMRI at T0 and T1. Second, we tested the *a priori* hypothesis based on (Saccaro, Delavari, Meuleman, *et al*., 2025; Saccaro *et al*., In Preparation) that MBCT-L increases coactivation between the hippocampus and the somatomotor network, quantified through hippocampus seed-based coactivation patterns (CAPs). We then examined, in exploratory analyses, whether the magnitude of this neural change scaled with the individual benefit participants derived from the intervention, a relationship that would position hippocampus-somatomotor coactivation as a candidate marker of treatment response.

## Material & Methods

### Participants

Eligibility criteria were applied to recruit second, third- or fourth-year healthcare students in Geneva, Switzerland, who were at least 18 years old, were not mindfulness experts, and agreed to refrain from regular mindfulness practice until allocated to the MBCT-L group. Eligible and willing students attended an in-person 1-hour visit with a psychologist at the Geneva University Hospital. At the end of the visit, the students were officially included in the study if corresponding to the eligibility criteria. Students were excluded if they had a current depressive episode, post-traumatic stress disorder, psychotic disorder, or substance dependency as identified through the Mini International Neuropsychiatric Interview (MINI) (Sheehan, Lecrubier et al. 1998). If not eligible and if needed, the students were referred to appropriate mental health specialists or the psychology consultations offered at the University of Geneva.

The study protocol was approved by the Geneva Ethics Committee (n°2021-02059). All participants received oral and written information about the trial and informed written consent was obtained from all participants before the beginning of the study. Out of the 161 healthcare students studying in Geneva University or Geneva School of Health Sciences (Geneva, Switzerland) enrolled in the study, 74 participants (54 females, 1 transgender person; age: 24 ± 5 (mean±sd) years old) agreed to perform the fMRI part of the study. From these 74 participants, 17 dropped out of the fMRI part of the study or of the study as a whole so that the sample of the current paper is constituted of 57 participants (39 female, 1 transgender person; age: 23 ± 5 years old). Participants were compensated for their participation with vouchers for a maximum total amount of 230 CHF if all steps were completed.

### Intervention

After inclusion, participants were randomly assigned to either the control group or the mindfulness intervention group (**Fig. 1**) through a dedicated online webpage managed by the Clinical Research Centre of the Geneva University Hospital. Participants and study staff were blinded during the screening visit before the randomization. The participants in the mindfulness intervention group followed the Mindfulness-Based Cognitive Therapy for Life (MBCT-L) guidelines (Bernard *et al*., 2017) which require to follow one pre-course orientation session, eight online sessions of 2h00 of MBCT-L in groups of up to 16 individuals over 8 weeks supervised by a MBCT-L trained instructor (3 instructors were involved in the study) and a half-day silent session of mindfulness also supervised by a MBCT-L instructor. All instructors had completed a full certified MBCT-L teacher training and had already led MBCT-L groups prior to the start of the study. Additionally, participants in the MBCT-L intervention group were also invited to practice meditation regularly at home and were provided with an access to a dedicated smartphone application developed at the University of Geneva to support mindfulness research across studies . To maximize the effects of the procedure, participants familiar with meditation practice were asked to refrain from regular mindfulness practice before being allocated to the MBCT-L group. All participants in the intervention group attended at least 6 of the 8 MBCT-L sessions during the experiment. Participants allocated to the control condition continued their usual activities and did not receive MBCT-L during the equivalent 8-week period.

**Figure 1:**
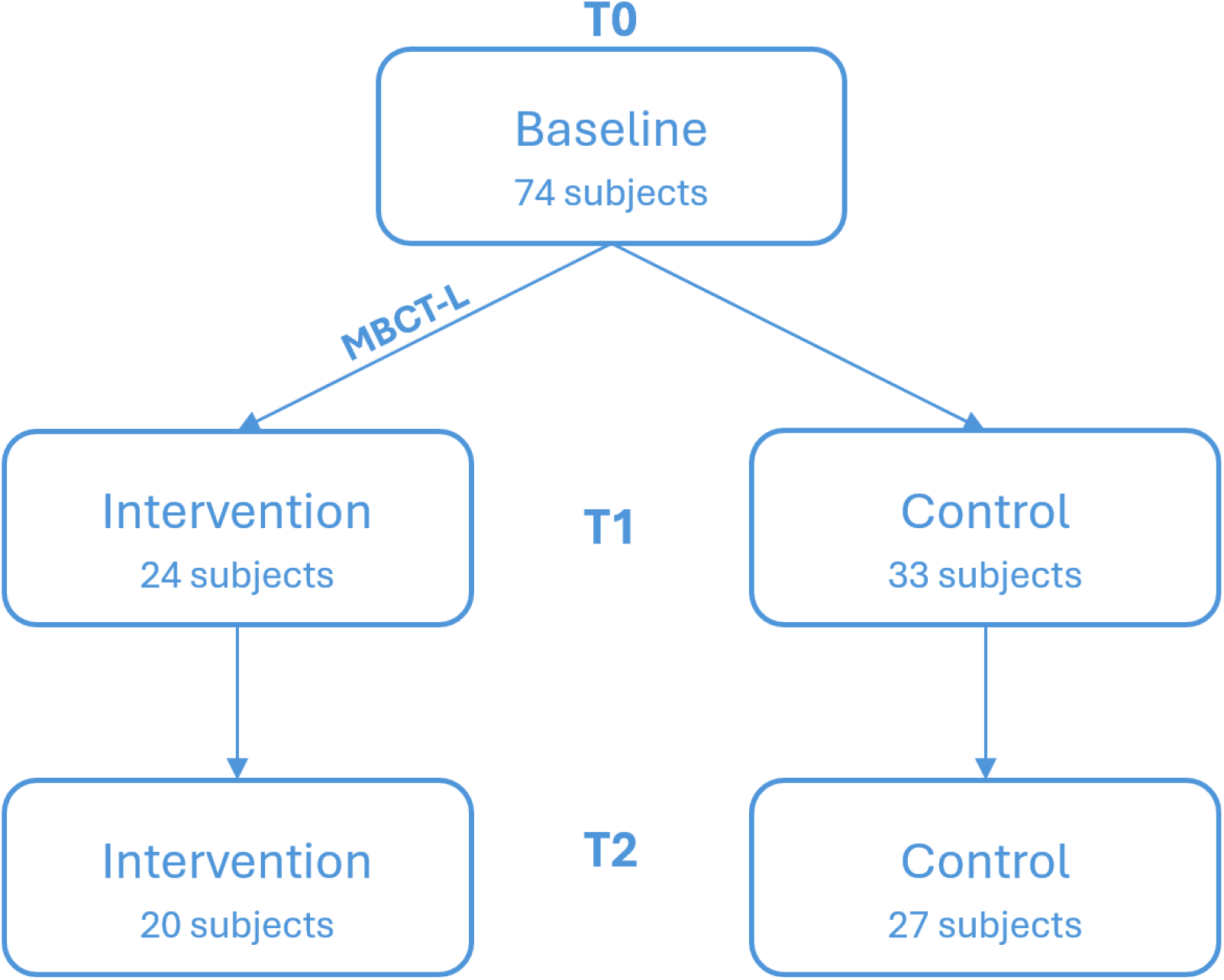
Experimental design. Out of the 161 individuals involved in the study, 74 were eligible and accepted to also perform the fMRI part (the rest only filled behavioral questionnaires without going into fMRI). They came for a first visit in the laboratory (T0). These subjects were then randomly assigned to either the control or the Mindfulness-Based Cognitive Therapy for Life (MBCT-L) intervention group. The individuals in the MBCT-L group performed eight 2h online sessions of MBCT-L supervised by a trained instructor, while the control group was unsupervised for a similar period of time. After this period, participants came for a second visit (T1). Finally, participants came for a third visit (T2) to assess the long-term effects of the intervention.

In each visit, participants filled out questionnaires out of fMRI and performed 8 minutes of resting-state fMRI followed by a T1-weighted anatomical scan. The participants also performed other behavioral tasks that will be developed in other publications and that did not affect the present results (see **Supplementary Material** for more information). The first visit T0 was performed at baseline before any intervention. After approximately 8 weeks, participants from both groups were invited to repeat the same procedure during the second visit T1. Finally, approximately 16 weeks after the start of the intervention (T0) and 8 weeks after its end (T1), participants were invited to come for the third visit T2 to assess the long-term positive effects of the intervention. Since several participants dropped out between visits T1 and T2, the data related to the third visit is only provided indicatively as we focused on T1 to maximize statistical power.

### Questionnaires

In each of the three visits, participants filled in several questionnaires aiming at 1) assessing whether MBCT-L improved their general capacity to enter in mindfulness states and 2) at exploring the impact of the MBCT-L intervention on other dimensions such as mental health and empathy (see **Supplementary Material** for more details).

The capacity to enter in mindfulness and the different aspects of mindfulness were assessed with the Five Facet Mindfulness Questionnaire (FFMQ) (Baer *et al*., 2006; Heeren *et al*., 2011) which output includes one global score and five subscores including 1) observing corresponding to the ability to attend to internal and external stimuli, 2) describing corresponding to the capacity to describe with words internal bodily states or emotions, 3) acting with awareness corresponding to the capacity to be attentive to one’s current actions, 4) non-judging inner experience corresponding to a non-evaluative stance toward inner thoughts and emotions , 5) non-reacting to inner experience, which corresponds to the ability to allow thoughts and emotions to flow without becoming preoccupied about them.

Additionally, on the second and third visits, the MBCT-L group only answered to a few additional questions regarding their own subjective assessment of how much the intervention was beneficial and led them to improve their stress handling, as well as a few supplementary questions related to their general wellbeing (sleep quality, concentration, exercise, use of drugs, taking time for one self, etc.) and their frequency of mindfulness practice on their own during the period of the intervention.

### fMRI data acquisition

Whole-brain MRI data were acquired with a 3T scanner (Siemens TIM Trio), and a 64 channels head-coil, at the Brain & Behavior Laboratory, University of Geneva. The Blood Oxygenation Level Dependent (BOLD) contrast was measured using a T2*-weighted echo-planar sequence (EPI). Resting-state fMRI data acquisition consisted in six hundred functional volumes of 54 slices each (TR = 720 ms, TE = 30 ms, flip angle = 52°, FOV = 210 mm, multiband SMS accelerating factor = 6, voxel resolution: 2.5*2.5*2.5 mm^3^). The subjects were instructed to stay awake, with normal breathing, and not think about anything for the 8 min duration of the scan. At the end of each visit, a high-resolution T1-weighted anatomical image was acquired with the following parameters: TR = 2000 ms, TE = 2.49 ms, flip angle = 9°, FOV = 230 mm, voxel resolution: 0.8*0.8*0.8 mm^3^, phase encoding direction: anterior–posterior.

### fMRI data preprocessing

Preprocessing of the fMRI data was performed with the SPM12 toolbox (Wellcome Trust Center for NeuroImaging) running in Matlab R2023a. Preprocessing consisted of spatial realignment, normalization using the same transformation as anatomic images, and spatial smoothing using a Gaussian kernel with a full width at a half-maximum (FWHM) of 5 mm. We then computed framewise displacement, a measure indexing head motion from one volume to the next based on the sum of the absolute values of the differentiated realignment estimates (by backwards differences) at every timepoint (Power *et al*., 2012; Power *et al*., 2014). Head movement analysis did not reveal major head movement, as there were only 6 volumes with framewise displacement head motion higher than 2mm and only 2 volumes with head motion higher than 3mm out of the 600 volumes/session*57 subjects*2 visits = 68 400 total volumes included in the analysis. Nevertheless, we scrubbed image volumes with frame-wise displacement (FD) above 0.5 mm. Additionally, six motion parameters, white matter, and cerebrospinal fluid (CSF) signals were regressed out from the data as nuisance variables to reduce the impact of nonneuronal BOLD fluctuations and motion on the signal.

### fMRI data analysis

The bilateral hippocampus seed has been described previously (Saccaro *et al*., 2023; Saccaro, Delavari, Van De Ville, *et al*., 2025; Saccaro, Delavari, Meuleman, *et al*., 2025; Saccaro *et al*., In Preparation), but further details can be found in the Supplementary Materials (**Supplementary Methods** and **Supplementary Figure 1**). The micro-coactivation patterns (μCAPs) extraction was performed with functions from the TbCAPs toolbox (https://github.com/MIPLabCH/TbCAPs; (Bolton *et al*., 2020)) running in Matlab R2023a. In brief, the μCAPs analysis consists in identifying the contributions of different subsections of the seed region to specific patterns of synchronous brain activity, known as coactivation patterns (CAPs). The TbCAPs toolbox extracts and Z-scores the seed BOLD time-series for each fMRI session, selecting the time points with the highest activity. Through K-means clustering algorithm, the common pool of the retained whole-brain volumes from the two visits (T0 baseline and T1 right after intervention) of all 57 subjects is classified in a data-driven manner into different activity-based clusters (CAPs), for which within-cluster differences (defined as spatial similarity) were smaller than across-cluster ones. The optimal number of micro co-activation patterns (K) was then determined using an automated elbow method on the within-cluster sum of squared errors (WCSS), selecting the K value that maximized the slope change ratio following iterative convergence of seed subdivisions (see **Supplementary Material** for more details). These individual CAPs are then converted into spatial Z-maps, to quantify the significance of their deviation from zero. The μCAPs analysis code is accessible at: https://github.com/MIPLabCH/mCAP and is comprehensively detailed elsewhere (Delavari *et al*., 2024).

We extracted for further analysis the temporal feature of CAPs’ occurrences, that is, defined as the sum of time points (frames) assigned to a given CAP among all the retained frames, across the entire timecourse. In other words, occurrences reflect the proportion of time that each subject’s CAP was co-active with the hippocampus seed.

The identification of each neural network was performed based on the correlation between the voxels of each CAP with the seven neural networks identified by Yeo and colleagues based on functional connectivity patterns (Yeo *et al*., 2011). Each CAP was then labelled according to its maximum degree of correlation.

### Pupil diameter

Pupil diameter was recorded at a sampling rate of 1 kHz, using an EyeLink 1000 Plus (SR Research) eye tracker during the whole 8 minutes of resting-state fMRI session. The pupil data was missing either for T0 or T1 in 8 participants due to recording issues and in one subject the fMRI triggers were not recorded forbidding to realign the data with the fMRI resting-state data so that the pupil analysis was focused on N = 49 subjects (N = 28 controls; N = 21 MBCT-L). Participants were asked to keep their eyes open during the whole session so the data was mostly usable. The eye tracker was calibrated before the start of fMRI sessions, once the subject was positioned inside the scanner. Any sample where the participant was looking outside of the screen, blinking or with a range further from [mean±3*SD] of the signal was removed and interpolated with MATLAB *interp1* function, which implements the pchip cubic interpolation method. Within-trial individual time series were then smoothed using a 150 ms kernel. Finally, since the distance between the camera and the eye was relatively constant across all participants and sessions, we used SR Research Light’s artificial eye procedure to determine the scaling factor that allows to convert the pupil diameter from arbitrary units to millimeters. Samples of a single visit where less than 20% of the 7.2 minutes of data acquisition were usable were completely discarded from the analysis leading to the exclusion of 28 more subjects as we only kept participants where data was usable in both visit T0 and T1. We therefore ran the analysis on the remaining 21 subjects (N = 9 controls; N = 12 MBCT-L).

### Heart-rate data acquisition

During each fMRI acquisition, participants were also wearing a photoplethysmograph (Biopac, USA) on the left index finger to measure heart rate and heart-rate variability during resting-state and task execution (sampling rate: 1 kHz).

### Heart-rate analysis

The signal was first filtered with a 0.5 to 3 Hz band-pass digital filter in Matlab. Then the intervals between heartbeats were estimated. Outliers were identified with Grubbs’s test and a shape-preserving piecewise cubic spline interpolation was performed in Matlab. We then extracted the average heart-rate, expressed in beats per minute (bpm), the Root Mean Square of Successive Differences between normal heartbeats (RMSSD) which is a measure of heart-rate variability (HRV) which mostly reflects parasympathetic vagal tone (Laborde *et al*., 2017; Shaffer and Ginsberg, 2017) and the pulse amplitude, whose fluctuations partly reflect (ortho)sympathetic tone (Njoum and Kyriacou, 2013; Budidha and Kyriacou, 2019). To ensure that the data made sense biologically, we also removed any heart-rate value outside of the [20 200] bpm range and any RMSSD value outside of the [10 120] ms range.

### Statistical analyses

All statistical analyses were performed in Matlab R2023a (The MathWorks). For most variables, unless explicitly stated otherwise, the main test performed was a repeated-measures ANOVA which aimed at assessing the Group*Time effect of the intervention. This tests the hypothesis that the MBCT-L intervention had an impact (Time: T1>T0) and whether this impact was greater in the MBCT-L group (Group: MBCT-L>Controls).

## Results

The main goal of the current study was to assess whether a MBCT-L intervention would induce brain activity changes at rest, particularly regarding the hippocampus-somatomotor coactivation pattern occurrence, reflecting changes in stress handling and the perceived impact of the intervention on the life of the participants. To do so, 57 participants came to the laboratory over two visits during which they filled behavioral questionnaires and performed resting-state functional magnetic resonance imaging (fMRI).

### MBCT-L is associated with higher self-reported mindfulness scores

Quite expectedly, the FFMQ total score, a questionnaire which reflects five different facets of mindfulness, displayed a significant and large Group*Time effect (**Supplementary Figure 2A**; F(1,55) = 5.776, p = 0.020, ηp^2^ = 0.095) confirming that the MBCT-L intervention induced improved mindfulness capacities. When looking at each of the five FFMQ subscores individually, we also observed that two of the five dimensions of the FFMQ were showing significant Group*Time effects including: the observing dimension (**Supplementary Figure 2B**; F(1,55) = 6.898, p = 0.011, ηp^2^ = 0.111), which reflects the capacity to observe one’s internal sensations and the non-reacting to inner experience dimension (**Supplementary Figure 2C**; F(1,55) = 4.820, p = 0.032, ηp^2^ = 0.081), which reflects the capacity to allow for the free flow of thoughts and emotions without being preoccupied about them. All effects reflect an increased score in the MBCT-L group after the intervention in comparison with the control group.

We also explored the other questionnaires, but except one significant Group*Time effect on the Acceptance and Action questionnaire (**Supplementary Figure 2D**; F(1,54) = 6.129, p = 0.017, ηp^2^ = 0.102, uncorrected for multiple comparisons) suggesting increased psychological flexibility after the MBCT-L intervention, there was no significant Group*Time effect on the other questionnaires that were assessed (all p > 0.05, uncorrected for multiple comparisons). This is also true for the Perceived Stress Scale (PSS) questionnaire (F(1,55) = 0.007, p = 0.930, ηp^2^ = 0.0001) which measures the degree to which individuals perceive situations in their lives as stressful, suggesting that there was no tremendous impact of the MBCT-L intervention on the perception of stress.

### Characterization of the brain coactivation patterns in the population

Based on a data-driven “consensus” procedure (Monti *et al*., 2003), six micro-coactivation patterns (μCAPs) co-activating with the hippocampus seeds were identified (**Figure 2A; Supplementary Figure 3**). Based on their overlap with known neural networks from the literature (Yeo *et al*., 2011), we observed that three of these CAPs are positively associated with the activation of the default mode network or DMN (CAP1, CAP2, CAP6), while the remaining CAPs were uniquely associated with distinct neural networks: frontoparietal network (CAP3-FPN), somatomotor network (CAP4-SMN), visual cortex (CAP5-V). The CAPs associated with DMN coactivation are clearly distinguished from each other by their negative associations with the limbic (CAP1-DMN_L_), dorsal attention network (CAP2-DMN_DAN_) or ventral attention network (CAP6-DMN_VAN_).

**Figure 2:**
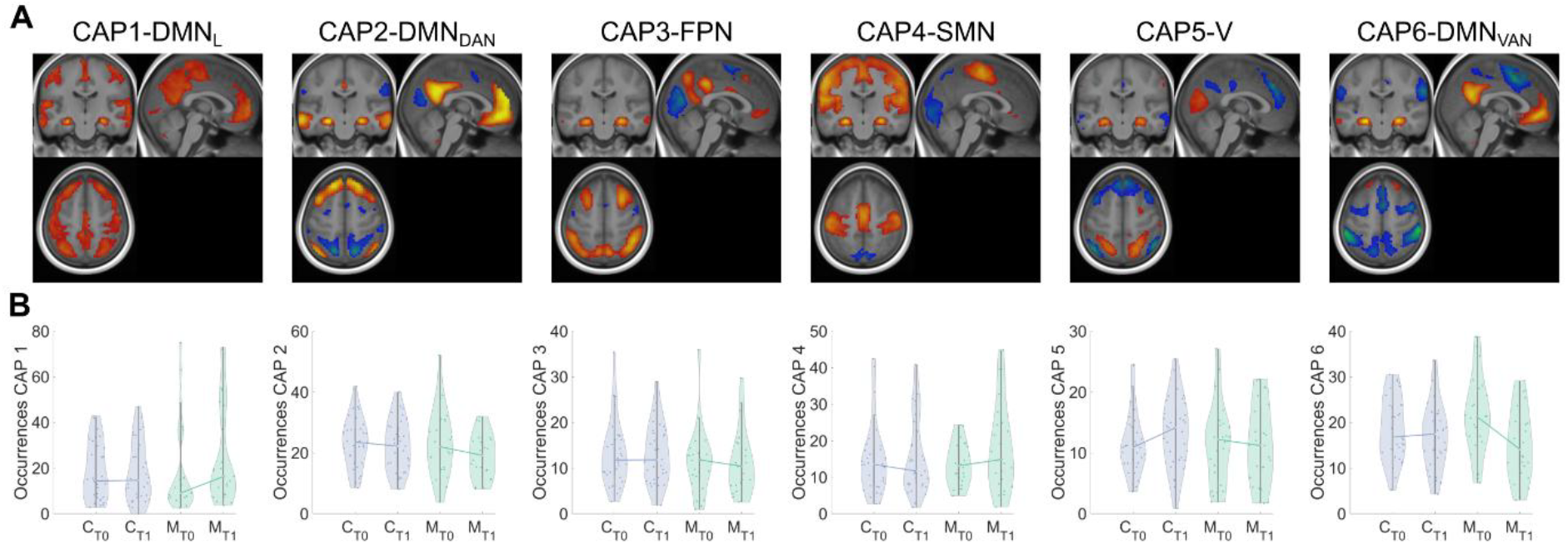
Hippocampus micro-coactivation patterns. A] Each image displays one hippocampus coactivation pattern (CAP) overlaid upon the average anatomical mask of the 57 participants over their 2 visits. For display purposes, each CAP has been thresholded so that only the top 25% of the voxels included in that CAP are visible. Each map is centered at the following MNI coordinates (x = 2; y = -20; z = 55) and displays both positive (with a red-to-yellow gradient) and negative (with a blue-to-green gradient) coactivation patterns. B] Violin plots highlighting the distribution of the CAP occurrences for each group (C: control group, displayed in blue; M: MBCT-L group, displayed in green) and each visit (T0: baseline; T1: after the intervention). The curves between T0 and T1 connect the median for each visit, to highlight the orientation of the changes between T0 and T1. There is no significant Group*Time effect for any of the CAPs.

### No significant global effect of the MBCT-L intervention on neuroimaging or physiology

Based on the aforementioned previous results from our group (Saccaro, Delavari, Meuleman, *et al*., 2025; Saccaro *et al*., In Preparation), we expected higher hippocampus-somatomotor network co-activity (CAP4-SMN) in the MBCT-L group after the intervention in comparison with the baseline and the control group (Group*Time effect). To assess this Group*Time effect, we performed a repeated-measures ANOVA which demonstrated that CAP4-SMN did not display a significant Group*Time effect (**Figure 2B**; F(1,53) = 1.60; p = 0.21, ηp^2^ = 0.03), even when repeating the same ANOVA while controlling for sex and age (F(1,51) = 0.87; p = 0.36, ηp^2^ = 0.02). However, there was still a small tendency in the expected direction of a MBCT-L-related enhancement of the hippocampus-somatomotor coactivation pattern occurrence (**Figure 2B)**. Notably, none of the other CAPs displayed a significant Group*Time effect (all p > 0.15, uncorrected for multiple comparisons) suggesting that there were no major group brain effects of the MBCT-L intervention. Given the strong expectation based on the literature that the DMN activity may decrease with MBCT-L, we nevertheless observed that, despite no significant Group*Time effect for the two CAPs associated with the DMN (CAP2-DMN_DAN_: F(1,54) = 2.253, p = 0.139, ηp^2^ = 0.04; CAP6-DMN_VAN_: F(1,55) = 1.026, p = 0.316, ηp^2^ = 0.02) as aforementioned which precludes from dawing any conclusion on this regard in our data, there was a moderate-to-large tendency for a DMN coactivation pattern occurrence decrease in the MBCT-L group for both DMN-related CAPs (CAP2-DMN_DAN_: T0: 23.297±2.232 > T1: 19.039±1.484, T1-T0: -4.26±2.14, d = 0.46; CAP6-DMN_VAN_: T0: 21.425±1.725 > T1: 15.182±1.567, T1-T0: -6.24±1.52, d=0.77) that was much smaller in the control group (CAP2-DMN_DAN_ T0: 23.240±1.425 > T1: 22.216±1.451, T1-T0: - 1.025±1.62, d=0.12; CAP6-DMN_VAN_ T0: 18.780±1.201 > T1: 17.176±1.233, T1-T0: - 1.60±1.34, d=0.23). Similarly, we did not observe any significant Group*Time effect in pupil diameter (**Supplementary Figure 4A**; F(1,21) = 0.180, p = 0.676, ηp^2^ = 0.009), heart-rate (**Supplementary Figure 4B**; F(1,54) = 0.065, p = 0.799, ηp^2^ = 0.001), heart-rate variability measured with RMSSD (**Supplementary Figure 4C**; F(1,48) = 0.216, p = 0.644, ηp^2^ = 0.005) or pulse amplitude (**Supplementary Figure 4D**; F(1,54) = 0.068, p = 0.796, ηp^2^ = 0.001) during the resting-state session.

### The hippocampus - SMN coactivation as a biomarker of MBCT-L impact

Despite the lack of significant brain and physiological changes with the MBCT-L intervention at the group level, inter-individual variability in CAP4-SMN activity may still reflect inter-individual variability in the effectiveness of the intervention, particularly regarding stress management and well-being. Therefore, we next focused on assessing whether higher increases in hippocampus - CAP4-SMN co-activity at rest after the MBCT-L intervention were associated with the subjective impact of the intervention and subjective stress handling. The results confirmed this hypothesis as there was a positive correlation between the increase in hippocampus - CAP4-SMN co-activity between T1 and T0 in the MBCT-L group and the subjective importance of the intervention measured at T1 (**Figure 3A**; r = 0.481, p = 0.017). This result was also confirmed by the correlation between the CAP4-SMN activity increase in the MBCT-L group and the subjective perception that the intervention had long-lasting benefits measured at T1 (**Figure 3B**; r = 0.412, p = 0.045). Furthermore, hippocampus - CAP4-SMN co-activity increases also tended to correlate with better stress handling after the intervention (**Figure 3C**; r = 0.352, p = 0.092). In contrast, there was no significant correlation between the CAP4 activity increase and the PSS score at T1 (r = -0.262, p = 0.215) or with the change of PSS score between T0 and T1 (r = 0.094, p = 0.663), signaling that CAP4 activity does not reflect any change in the perception of how stressful situations are. Furthermore, the correlation with long-lasting benefits tended to remain stable at T2 (**Figure 3D;** long-lasting benefits: r = 0.531, p = 0.011), which means that the heightened hippocampus-CAP4 activity at T1 is not only associated with improved well-being and stress handling at the end of the intervention but is also predictive of the subjective benefits of the intervention approximately two months after the end of the intervention.

**Figure 3:**
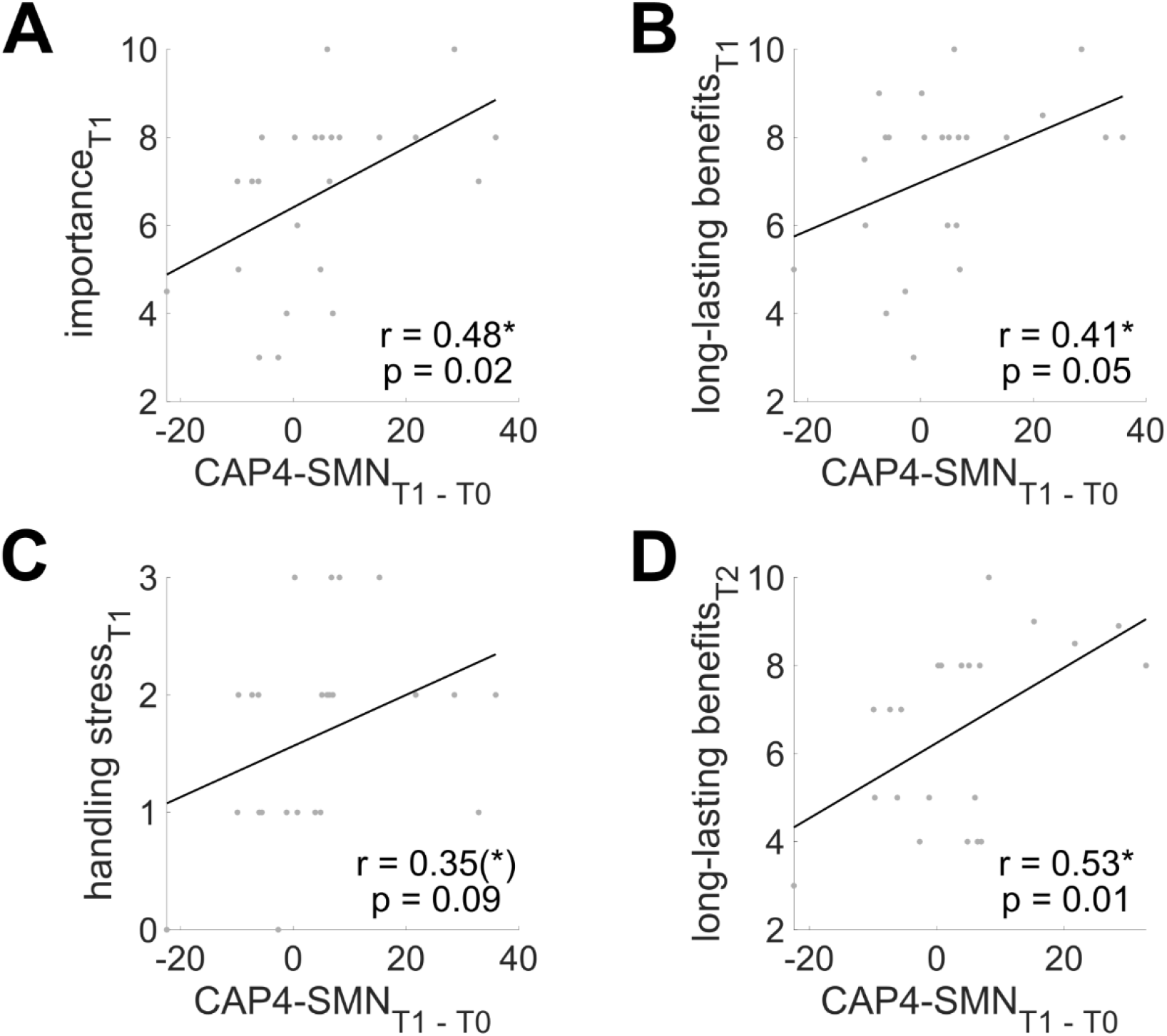
Correlation between the hippocampus – CAP4-SMN activity increase and the subjective quality of the intervention in the MBCT-L group. Correlation between the increase of the hippocampus – CAP4-SMN coactivation pattern between T0 and T1 and A] the subjective importance of the program measured at T1; B] the subjective long-lasting benefits of the intervention measured at T1; C] the subjective capacity to handle stress at T1 and D] the subjective long-lasting benefits of the intervention measured at T2. The analysis is restricted to the participants of the MBCT-L group (N = 24 at T1 for panels A to C; N = 22 at T2 for panel D). Significance of the Pearson’s correlation: *p < 0.05; **p < 0.01; ***p < 0.001.

We next asked whether the magnitude of the CAP4-SMN increase scaled with the change in self-reported mindfulness over the intervention, as assessed with the FFMQ questionnaire. Within the MBCT-L group, the change in total FFMQ score from T0 to T1 was not significantly associated with the CAP4 increase (**Figure 4A**; r = 0.261, p = 0.217), indicating that the relationship was not tied to global mindfulness improvement. However, examining the five FFMQ facets independently, the change in non-reactivity (the capacity to let thoughts and emotions arise and subside without being caught up in them) correlated with the CAP4-SMN increase (**Figure 4F**; r = 0.491, p = 0.015), with a weaker trend for non-judging of inner experience (the tendency to take a non-evaluative stance toward one’s thoughts and feelings; **Figure 4E**; r = 0.383, p = 0.065). No other facet showed any relationship (**Figure 4B-D**; all p > 0.4). Given the number of facets and time windows examined, these associations are reported without correction for multiple comparisons and should be considered exploratory. With that caveat, the pattern suggests that the MBCT-L intervention increase in hippocampus-related somatomotor coactivation tracks improvement in a specific affect-regulatory facet of mindfulness which reflects emotion regulation (observing internal experience without reacting to it) rather than a global change in mindfulness.

**Figure 4:**
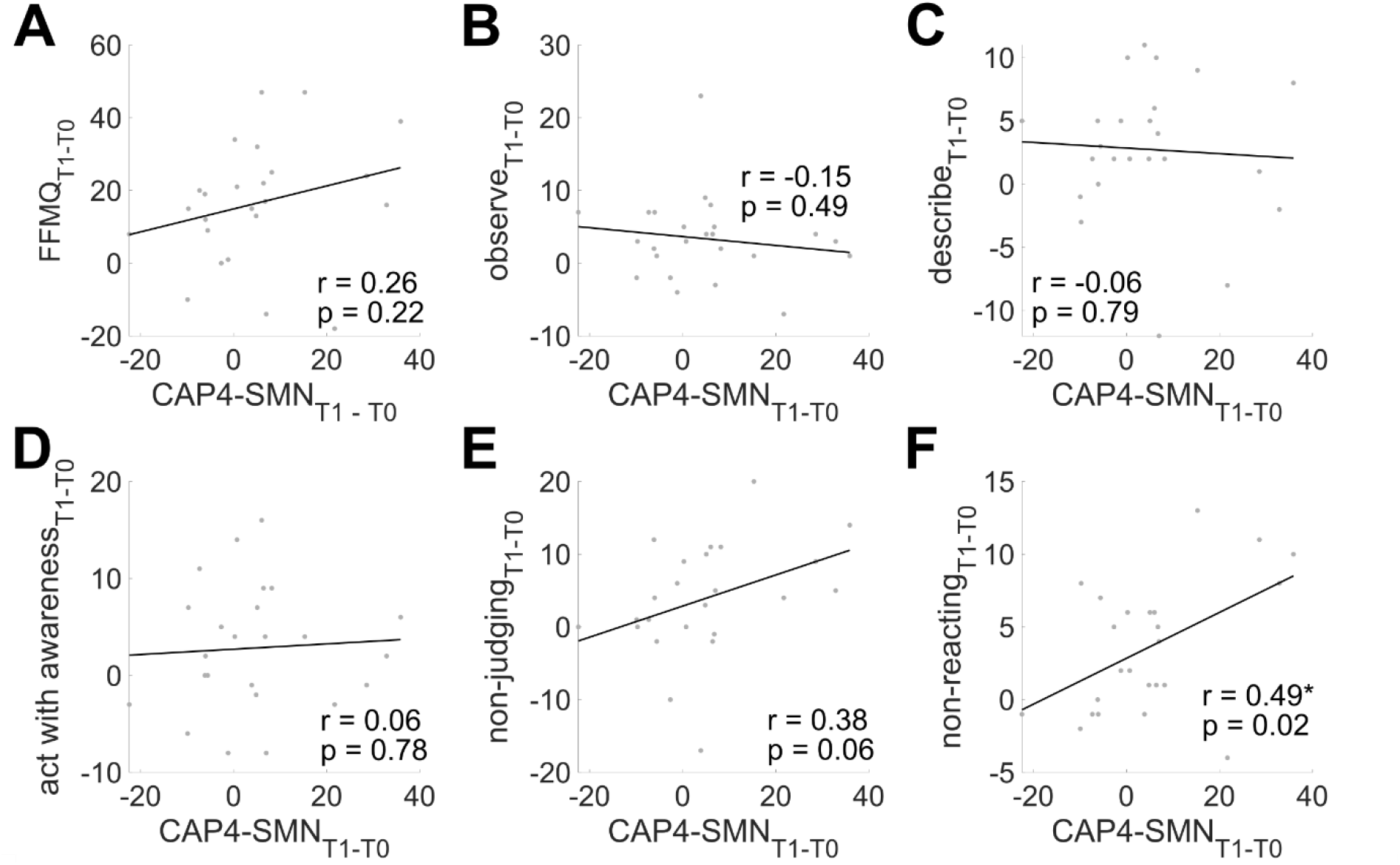
FFMQ scores in relationship with the hippocampus - CAP4-SMN activity changes between T0 and T1. A] FFMQ total score change between T0 and T1 in function of the change in CAP4-SMN occurrences between T0 and T1. B-F] FFMQ subscores change between T0 and T1 in function of the change in CAP4-SMN occurrences between T0 and T1 following the order: B] observe, C] describe, D] act with awareness, E] non-judging and F] non-reacting subscores. Significance of Pearson’s correlation: (*)p < 0.1; *p < 0.05; **p < 0.01; ***p < 0.001.

## Discussion

In this randomized controlled trial in 57 healthcare students, an 8-week MBCT-L intervention produced a reliable improvement in self-reported mindfulness, confirming that the intervention engaged its primary psychological target. Within the FFMQ subscores, the strongest Group*Time effects concerned the observing and the non-reactivity to inner experience subscores. Notably, these results, focused on the 57 participants that underwent resting-state fMRI, are also replicated in the analyses that include all the participants of the behavioral study (Jermann *et al*., In Preparation), confirming that the MBCT-L was effective in driving mindfulness and that the results observed here are not specific to the subsample of participants we focused upon. While the global score reflects the effectiveness of the intervention, the observing and non-reactivity to inner experience dimensions point towards improved stress management and emotion regulation after the intervention, as they reflect the ability to notice internal experiences and allow thoughts and emotions to arise without immediately reacting to them. In parallel, the absence of any significant difference in the perceived stress scale after the intervention signals that individuals did not experience less frequently stressful situations in their daily lives but were maybe better equipped to deal with them after the intervention. This pattern suggests that MBCT-L may primarily strengthen metacognitive and decentering-related capacities leading to improved emotion regulation, rather than simply reducing perceived stress exposure.

At the level of resting-state brain dynamics and physiology, the picture was more nuanced. While we replicate the observation that hippocampus micro-coactivation patterns during resting-state can be summarized with 6 main coactivation patterns (Saccaro *et al*., In Preparation), we did not detect a significant Group*Time effect of the intervention on hippocampus-related micro-coactivation patterns, nor on any physiological variable. However, rather than treating this dissociation as a null result to be dismissed, we consider it as a central observation of the study: in a non-clinical population, the effects of MBCT-L may not be uniform across all participants but instead depend on their baseline state at the onset of the intervention.

More specifically, exploratory analyses within the MBCT-L group showed that individuals with greater increases in hippocampus-SMN occurrences reported greater perceived benefits from the intervention, greater perceived long-lasting effects, and higher post-intervention non-reactivity to inner experience mindfulness scores. These findings suggest that hippocampus-SMN coactivation may serve as a biomarker indexing inter-individual variability in improved emotion regulation. Previous evidence has related the

SMN activity with emotion regulation, and particularly with the capacity to accept one’s emotions (Jiang *et al*., 2026), in line with our own observation that the MBCT-L-related increased hippocampus-SMN activation correlated with increased non-reactivity and non-judging of one’s internal emotions. Previous research from our own research group also shows that the hippocampus-SMN coactivation pattern occurrence is a transdiagnostic biomarker of emotion dysregulation disorders, even in the patients’ offspring which are at risk of developing emotion dysregulation disorders (Saccaro *et al*., 2023; Saccaro, Delavari, Van De Ville, *et al*., 2025; Saccaro, Delavari, Meuleman, *et al*., 2025). Additionally, our research in another sample also demonstrates that a MBCT-L intervention in teenagers increases the activity of this network (Saccaro *et al*., In Preparation). The present results suggest that the capacity to increase the activity of this hippocampus-SMN network by mindfulness practice leads to an improved capacity to accept one’s emotions and to handle stress and accounts, at least in part, for the subjective benefits of mindfulness practice.

The exploratory decreases in coactivation pattern occurrence within the MBCT-L group of the hippocampus-DMN CAPs related to the ventral and dorsal attention networks (CAP2-DMN_DAN_ and CAP6-DMN_VAN_) are also consistent with previous mindfulness literature reporting reduced DMN activity at rest in long-term meditators (Berkovich-Ohana *et al*., 2016). This might be driven by the fact that mindfulness practice reduces DMN activation as compared to rest (Ganesan *et al*., 2023) so that regular practice may also impact resting-state brain activity. However, because the effects we observed were not supported by significant Group*Time interactions and were restricted to hippocampus coactivation patterns, they should be taken with caution. They only suggest that hippocampus-related DMN coactivation may warrant further investigation in larger samples.

Finally, our stress-handling results, though borderline and in need of replication, are coherent with the broader interpretation of the hippocampal-SMN network as a biomarker of mindfulness’ impact. Indeed, although the MBCT-L intervention did not change the perception of how stressful events are as there was no Group*Time effect, nor any significant correlation between the hippocampus-SMN activity increase and the PSS scale which measures “the degree to which situations in one’s life are appraised as stressful” (Cohen *et al*., 1983), we did observe that the hippocampus–SMN coactivation increase was associated, although non significantly, with better subjective stress handling, such that participants in whom the intervention had a greater effect reported feeling more capable of managing stressful situations. In other words, based on our results, the participants who underwent the MBCT-L intervention, and those most affected by it in terms of hippocampus-SMN coactivation increase, do not report experiencing fewer stressful events in their lives, but may rather feel more capable of handling such situations. This result is confirmed by with other studies observing increased emotion regulation after mindfulness interventions (Goldin and Gross, 2010; Britton *et al*., 2012; Wenzel *et al*., 2021).

### Limitations

The current study has several limitations. Foremost, the brain findings that motivate our central interpretation are within-group correlations in the MBCT-L arm alone (n = 24). Because we did not test whether the same brain–behavior coupling is present in the control arm, and because the group-level Group*Time effect on CAP4-SMN was not significant, these associations cannot be attributed definitively to the MBCT-L intervention; an equivalent coupling in controls would indicate that the relationship reflects stable individual differences rather than intervention-related change. The associations are additionally uncorrected for the number of facets and analyses examined and would not survive standard multiple-comparisons correction, so they must be considered as exploratory and hypothesis-generating throughout.

Second, the trial was unblinded and used a passive control (no-intervention) which limits also the strength of the results as regular online meetings may bear a psychological effect on their own that cannot be assessed with a purely passive control. Additionally, every outcome that improved at the group level was self-reported (mindfulness, perceived benefit, subjective stress handling), whereas the objective resting-state and physiological measures showed no Group*Time effect, a pattern that expectancy effects are sufficient to produce and that the present design cannot exclude. An active comparator matched for attention, group contact and expectation would be needed to isolate effects specific to mindfulness in future studies.

Third, although the study as a whole involved a high amount of healthcare students (N = 161) over a relatively small data acquisition period (Spring 2022 - Fall 2023), the neuroimaging subsample used in the present study was relatively limited (N = 57 and only 24 in the MBCT-L group at T1). Since the effects of mindfulness on stress, well-being and physiological outcomes in non-clinical samples are expected to be small-to-moderate (Goyal *et al*., 2014; Kunzler *et al*., 2020; Goldberg *et al*., 2022), this remains limitation to be able to observe effects of MBCT-L in a non-clinical adult population. To confirm the robustness of these findings, future research should aim to replicate the observed effects with more substantial sample sizes.

## Conclusions and perspectives

In conclusion, MBCT-L improved self-reported mindfulness in healthcare students but did not induce a significant group-level increase in hippocampus-SMN CAP occurrence or resting physiological change. Nevertheless, despite its limitations, our study, along with other research highlighting the role of this brain network, provides key elements suggesting that the hippocampus-SMN coactivation is a promising biomarker of mindfulness’s impact on emotion regulation and stress handling. Indeed, exploratory individual-difference analyses suggest that participants who showed greater increases in hippocampus-SMN coactivation also reported greater subjective benefit, higher mindfulness and better stress handling after the intervention. These findings support hippocampus-SMN coactivation as a candidate neural correlate of individual benefit from mindfulness, particularly in relation to embodied and non-reactive emotion regulation processes. Replication in larger, adequately controlled studies, is required before this measure can be considered a good biomarker of mindfulness response but our study provides promising elements on this regard. Additionally, causal manipulations of this network, e.g. with repetitive transcranial magnetic stimulation of the SMN in patients with emotion dysregulation could also help to identify the involvement of this network on emotion regulation.

## Data availability

The data that support the findings of this study are available from the corresponding author upon reasonable request.

## Clinical Trial Registration

This trial was registered at ClinicalTrials.gov (NCT07110792).

## Funding

The project was funded by a grant from the Swiss National Science Foundation (SNSF) Grant n°32003B_205111/1.

## Conflicts of Interest

The authors declare no conflicts of interest.

## Notes

### Competing Interest Statement

The authors have declared no competing interest.

### Summary of Updates

- remove a typo by which one part of a sentence was in red - add Clinical Trial number that was missing + data availability, funding and conflict of interest sections that were missing - add Claudia Cachet's ORCID ID

