## Supplementary Material for "Hippocampal-somatomotor coactivation and perceived benefit following Mindfulness-Based Cognitive Therapy for Life in healthcare students: An exploratory randomized study"

### Supplementary Material & Methods

#### Questionnaires

In each of the three visits, participants filled several questionnaires aiming at measuring their mental health, their empathy for themselves and for others, and their general capacity of entering into mindfulness.

Mental health was assessed by measuring their levels of stress, anxiety, depressive symptoms, or intrusive thoughts with questionnaires in French including the Perceived Stress Scale (PSS) (Cohen *et al.*, 1983; Lesage *et al.*, 2012), the Hospital Anxiety and Depression Scale (HADS) (Zigmond and Snaith, 1983; Lépine *et al.*, 1985), the Psychological Well-Being Scale (PWBS) (Ryff and Keyes, 1995), the Sleep Condition Indicator (SCI) (Espie *et al.*, 2014; Bayard *et al.*, 2017), the Brief Resilience Scale (BRS) (Smith *et al.*, 2008; Jacobs and Horsch, 2019), the Perseverative Thinking Questionnaire (PTQ) (Ehring *et al.*, 2011; Devynck *et al.*, 2017), the Cognitive Fusion Questionnaire (CFQ) (Gillanders *et al.*, 2014; Dionne *et al.*, 2016), the Adult attention deficit/hyperactivity disorder Self-Report screening scale for DSM-5 (ASRS-5) (Ustun *et al.*, 2017; Baggio *et al.*, 2021), the Acceptance and Action Questionnaire-II (AAQ2) (Hayes *et al.*, 2004; Monestès *et al.*, 2009) and the Scale of Satisfaction with Studies (SSS).

Empathy for themselves and for others was assessed with the Prosocial Behavior Scale (PBS) (Caprara *et al.*, 2005; Carrizales *et al.*, 2019) and the Sussex-Oxford Compassion Scale for the self (SOCSS) and for others (SOCSO) (Gu *et al.*, 2020).

The capacity to enter in mindfulness and the different aspects of mindfulness were assessed with the Five Facet Mindfulness Questionnaire (FFMQ) (Baer *et al.*, 2006; Heeren *et al.*, 2011) which output includes one global score and five subscores including 1) observing corresponding to the ability to attend to internal and external stimuli, 2) describing corresponding to the capacity to describe with words internal bodily states or emotions, 3) acting with awareness corresponding to the capacity to be attentive to one's current actions, 4) non-judging inner experience corresponding to a non-evaluative stance toward inner thoughts and emotions , 5) non-reacting to inner experience, which corresponds to the ability to allow thoughts and emotions to flow without becoming preoccupied about them.

Additionally, on the second and third visits, the mindfulness group only answered to a few additional questions regarding their own subjective assessment of how much the intervention was beneficial and led them to improve their stress handling, as well as a few supplementary questions related to their general wellbeing (sleep quality, concentration, exercise, use of drugs, taking time for one self, etc.) and their frequency of mindfulness practice on their own during the period of the intervention.

#### Experimental design

During each visit, participants filled questionnaires, performed one resting state fMRI session and then two behavioral tasks including an adapted version of the Montréal Imaging Stress Task (Celen *et al.*, 2024) and the Socio-affective Video Task (Klimecki *et al.*, 2013). Importantly, the behavioral tasks were always performed after the acquisition of the resting-state so that the resting-state results were not influenced by the tasks.

### fMRI data analysis

#### *Bilateral Hippocampus seed*

The bilateral hippocampal seed was the same as in previous publications where the details of the way it was delimited can be found more extensively (Saccaro *et al.*, 2023; Saccaro, Delavari, Van De Ville, *et al.*, 2025; Saccaro, Delavari, Meuleman, *et al.*, 2025; Saccaro *et al.*, In Preparation) and it can be observed in **Supplementary Figure 1**. In brief, the seed aims to identify the hippocampus based on a combination of structural and functional parcellations.

#### *Co-activation patterns identification*

To identify the optimal number of micro-coactivation patterns (μCAPs), we tested a range of candidate cluster numbers (*K*) from 2 to 10. For each *K*, an iterative clustering procedure was performed to ensure stable seed parcellation. The functional data were divided into training and testing subsets. K-means clustering was applied to the training frames exhibiting significant seed activity (thresholded at $z > 0.5$), using a correlation distance metric, 100 replicates, and a maximum of 300 iterations per replicate. The clustering process was iterative: the resulting cluster centroids were used to update the seed mask subdivisions for the subsequent iteration. Convergence was assessed at each step by computing the spatial distance between the seed subdivisions of consecutive iterations. Specifically, the optimal matching between subdivisions was determined using the Hungarian algorithm (Munkres algorithm) on the pairwise cosine distance matrix. The procedure was considered converged when the mean distance between matched subdivisions fell below 0.005, or after a maximum of 8 iterations.

The optimal *K* was selected using an automated elbow method based on the stability of the clustering solution. For each tested *K*, we computed the mean of the maximum sum of distances. The optimal *K* was defined as the value maximizing the ratio of slope change: $\text{Ratio}_K = \left|\frac{\text{WCSS}_K - \text{WCSS}_{K-1}}{\text{WCSS}_K - \text{WCSS}_{K+1}}\right|$ or, in other words, as the inflection point (the "elbow") of this curve (**Supplementary Figure 3**) where the rate of decrease in the distance metric sharply decelerates indicating that additional clusters yield diminishing returns in terms of spatial variance explanation. This criterion identifies the point of maximal inflection on the WCSS curve, balancing model fit and parsimony without relying on visual inspection.

### Supplementary Figures

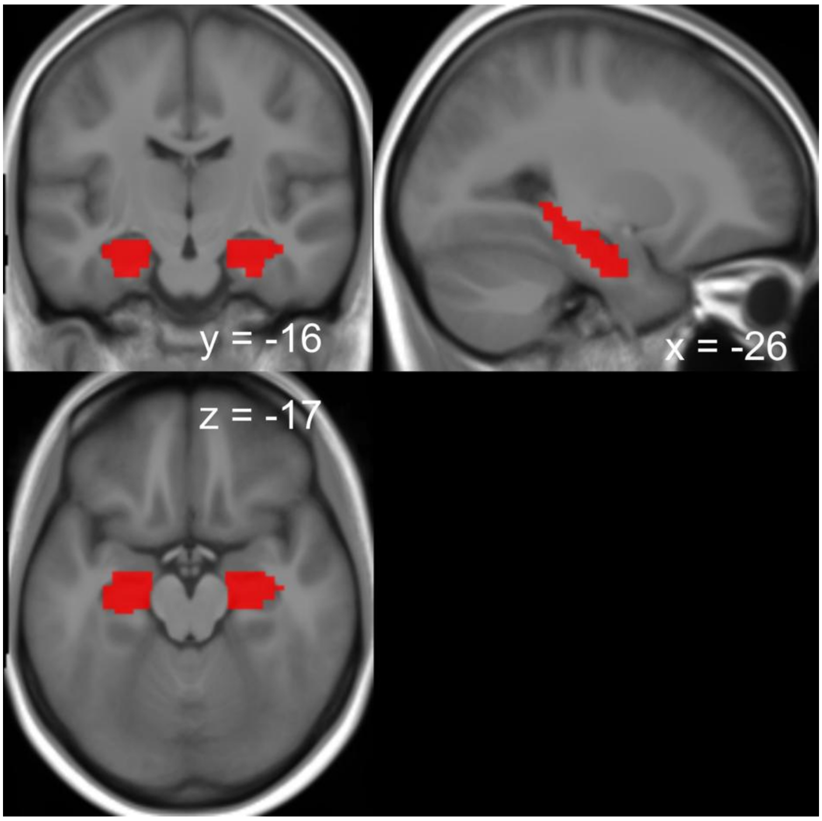


**Supplementary Figure 1: Hippocampal seed for micro-CAP analysis.**

The bilateral hippocampal seed is displayed overlaid on an averaged image of the anatomical brain of the 57 participants of the study across the visits T0 and T1. The MNI coordinates corresponding to each slice are displayed on the figure.

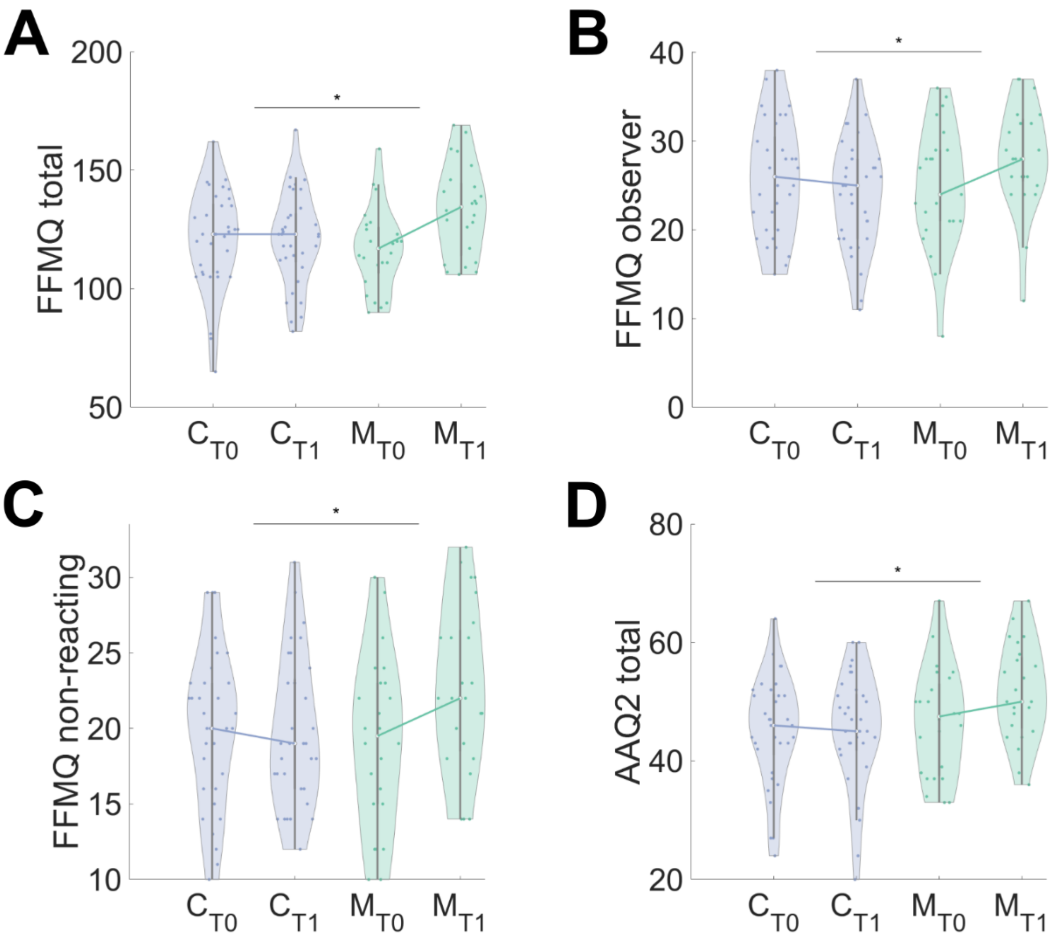


**Supplementary Figure 2: Questionnaires that display a significant Group*Time effect.**

A] FFMQ total. B] FFMQ observe score. C] FFMQ non-reacting to inner experience score. D] AAQ score. Significance of the Group*Time effect in a repeated measures ANOVA: *$p < 0.05$; **$p < 0.01$; ***$p < 0.001$ (uncorrected for multiple comparisons). The abbreviations indicate each group (C: controls in blue; M: MBCT-L in green) and each visit (T0/T1).

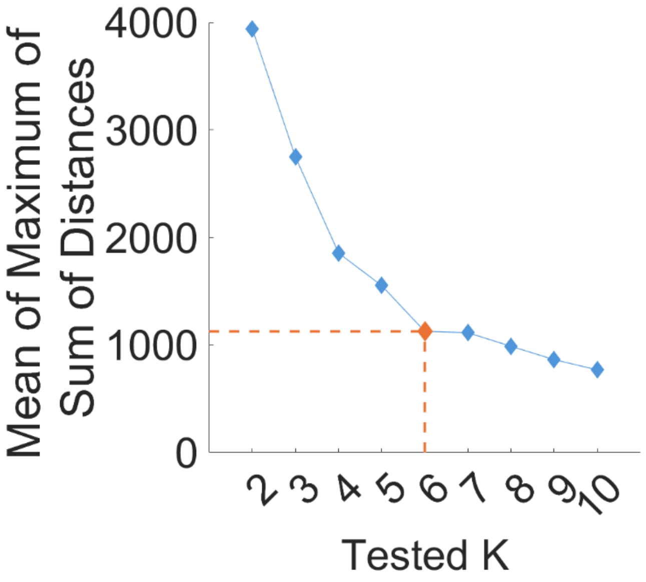


**Supplementary Figure 3: Determination of the optimal number of CAPs co-activating with the hippocampus across the whole population.**

The orange dotted line indicates the elbow of the curve corresponding to the optimal number of *K* co-activation patterns for the current dataset which includes the visits T0 and T1 of the 57 participants.

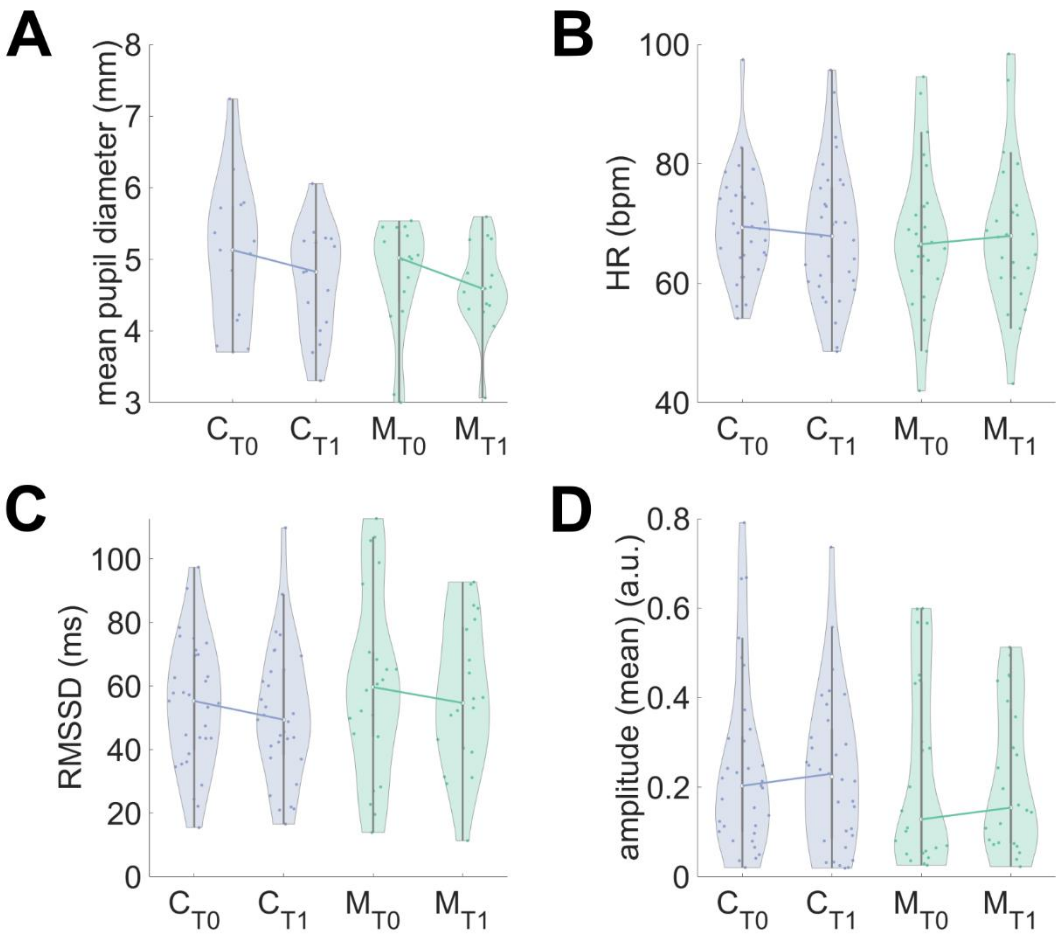


**Supplementary Figure 4: Physiological variables across groups and visits.**

A] Mean pupil diameter, B] Heart rate, C] RMSSD and D] pulse amplitude for each subject split by group and visit. The Group*Time effect was not significant ($p > 0.05$) in any of the variables tested. The curves link the medians between the two visits. The abbreviations indicate each group (C: controls in blue; M: MBCT-L in green) and each visit (T0/T1).
